# Peer victimization in adolescence alters gene expression and cytokine profiles during transition to adulthood

**DOI:** 10.1101/2024.07.24.604965

**Authors:** Jens Heumann, Steve W. Cole, Manuel Eisner, Denis Ribeaud, Edna Grünblatt, Michael J. Shanahan

## Abstract

Social adversity downregulates antiviral genes and upregulates inflammatory genes, altering cytokine levels. This study investigated the effects of peer victimization (PV) during adolescence on gene expression, cellular dynamics, and cytokine profiles in young adulthood. Data from the Zurich Brain and Immune Gene Study (z-GIG, *n* = 200; 47% female) captured PV between ages 11 and 20, with mRNA and plasma cytokines analyzed in 144 participants at age 22. A counterfactual design with genetically informed inverse probability weighting adjusted for baseline confounders in individual and environmental characteristics. *In silico* deconvolution revealed altered leukocyte composition in victims, including an M2-like monocyte–skewed profile. Differentially expressed genes in victims were enriched in preselected Reactome pathways, including *Interferon signaling, Metabolism* and *Signal transduction*. Exploratory Reactome-wide queries confirmed pathways in the *Immune System* and *Metabolism*, highlighted their downregulation, and identified additional pathways, such as those involved in *Chromatin organization* and *Metabolism of proteins*. Transcription factors STAT2, IRF2, and NF-κB emerged as key regulators in victims, with target genes prominently mapped to Interferon signaling and Chromatin organization pathways. Victims exhibited altered cytokine levels, including pro-inflammatory CCL4, TNF, CXCL9, and CXCL10. These findings suggest that adolescent PV can lead to changes in gene regulation and cytokine levels, potentially altering the immune profile into young adulthood. The results underscore the importance of health policies aimed at prevention, building resilience, and mitigating the long-term effects of PV.

## 1. Introduction

Peer victimization (PV) among adolescents, characterized by sometimes-traumatic experiences of verbal and physical abuse as well as systematic social exclusion, not only defines a challenging youth experience but also entails pronounced stress and may contribute to stress-related pathology, with effects similar to those observed in situations of forceful social subordination (Sapolsky, 2004). Such stress poses long-term consequences, including psychological distress like anxiety, depression, agoraphobia, externalizing problems, general anxiety disorder, panic disorder, psychosis, substance use, and self-harm (Klomek et al., 2015; Lereya et al., 2015; Moore et al., 2017; Reijntjes et al., 2011, 2010), and may even lead to suicide (Cha et al., 2018; Koyanagi et al., 2019; Van Geel et al., 2022). Moreover, PV is linked to physical stress symptoms, including headaches, asthma, skin problems, poor appetite, abdominal pain, vomiting, stiffness, backaches, dizziness, sleeping difficulties, and fatigue (Gini and Pozzoli, 2009, 2013; Hager and Leadbeater, 2016). Given the high global prevalence of PV, estimated at around 30% (Biswas et al., 2020), and its profound implications, a deeper understanding of the complex molecular impacts of PV could inform public health initiatives.

The occurrence of PV as a potent social stressor during adolescence introduces developmental considerations, as this period is marked by significant, possibly permanent, changes in the neuroendocrine, transcriptional, and immune systems that are critical for stress regulation (Casey et al., 2008; Rowson et al., 2019). These changes can in turn influence cytokine regulation (Chiang et al., 2012; Giletta et al., 2018; Polacchini et al., 2018), which subsequently feeds back into neuroendocrine processes (Dantzer et al., 2008) and creates a “neuroimmune pipeline” with implications for future health (Miller and Cole, 2012).

Biologically-informed PV research has focused on individual differences in the stress response, such as hair cortisol levels (e.g., Brendgen et al., 2023; Ouellet-Morin et al., 2021), cytokine activity (e.g., Chiang et al., 2012; Copeland et al., 2014; Giletta et al., 2018), epigenetic aging (e.g., Perret et al., 2023; Shalev et al., 2013), and DNA-methylation and other epigenetic processes (e.g., Marzi et al., 2018). While one study investigated immunomodulatory gene expression in homophobic victimization (Li et al., 2020), no such analysis has been conducted for PV. Studying gene expression is crucial for understanding how PV affects health, however, because it reveals the underlying biological mechanisms that link this common, salient stress to modulation of the immune system.

Furthermore, previous studies were limited by analyzing the biomolecular outcomes separately in different samples, by lacking longitudinal data on PV, and by inability to address potentially influential, time-varying socio-environmental confounders (Zuber et al., 2023). Findings on the impact of PV may also be biased by common-method variance, as many studies rely on self-reported measures for both PV and health-related outcomes (Gini and Pozzoli, 2013). Additionally, as PV studies are observational for ethical reasons, their inherent susceptibility to treatment selection bias should be acknowledged and addressed (Schoeler et al., 2018).

Social adversity is often associated with a characteristic transcriptomic profile, shaped by evolutionary adaptation, and characterized by upregulated pro-inflammatory genes and downregulated Type I interferon (antiviral) genes—a pattern referred to as the “conserved transcriptional response to adversity” (CTRA; Cole, 2019). Chronic activation of the CTRA is associated with an increased risk of psychological symptoms such as depression and anxiety, inflammation-related diseases such as cardiovascular diseases, metabolic diseases, neoplastic diseases and cancer, and downregulated immune function such as viral infections and a reduced response to vaccines (Cole, 2019, 2013). Research into the CTRA profile and PV remains nascent. Furthermore, studies reveal significant variations in transcription factor (TF) activity due to low SES—a social adversity with parallels to PV in terms of stress—with, for example, higher NF-κB and CREB activity, and lower GR (and consequently higher NF-κB:GR ratio) binding motif abundance (Miller et al., 2009; Ravi et al., 2024). This finding warrants the inclusion of TF activity analysis in genomic studies of PV.

In this study, 22-year-olds who experienced PV (victims) between ages 11 and 20 were contrasted with unaffected controls using a longitudinal design incorporating time-varying confounders. Leukocyte composition was estimated via *in silico* cell-type deconvolution (Avila Cobos et al., 2020). Comparisons were made for CTRA profile regulation and enrichment in preselected functional Reactome pathways, including immune-related functions, derived from a genome-wide analysis of differentially expressed (DE) genes. Additionally, an exploratory Reactome-wide query identified further pathways. TF activity and target genes were examined based on TF binding sites of DE genes. Finally, cytokine data, including 16 pro-inflammatory and other cytokines measured via a Multiplex immunoassay, were analyzed.

To address potential selection bias, genetically-informed inverse probability weighting (IPW) was applied across all analyses. The model incorporated pre-victimization characteristics, polygenic risk scores (PRS) for attention-deficit/hyperactivity disorder (ADHD), general anxiety disorder (GAD), major depressive disorder (MDD), and panic disorder (PD), as well as principal components from genotyping to account for underlying population structure.

## 2. Materials and Methods

### 2.1. Sample Characteristics

Data come from the Zurich Brain and Immune Gene Study (z-GIG). z-GIG is a subsample from z-proso (Ribeaud et al., 2022; z-proso Project Team, 2024), a longitudinal panel (n ≈ 1,500) started in 2004 with Zurich elementary school children aged 7–9, notable for high ethnic heterogeneity (> 60%) of the parents born abroad in 90 countries). In 2018, when eight waves had been completed (average age 20.5 years), for the z-GIG study, participants in z-proso were stratified based on self-reported peer adversity as well as relevant characteristics such as sex, teacher-rated peer adversity, aggression level, anxiety level, and conflict resolution skills. Individuals were assigned to the same stratum if they had the same sex and fell into the same median split on each of these variables (see Table S1 for the full list of variables and details of the 23 strata used for sample selection). 500 eligible subjects were randomly drawn from 23 strata and 200 finally participated in z-GIG at age 22 (response rate 40%), of whom 186 consented to blood sampling. No evidence of systematic nonparticipation was detected. IPW addressed selection bias from sample stratification and baseline confounders (explained *infra*, 2.4.). Valid mRNA and cytokine samples were obtained from 153 z-GIG participants. After one exclusion due to pregnancy and eight exclusions for statistical reasons related to IPW, the analytic sample comprised 144 participants, including 16 (11.1%) victims of PV (explained *infra*, 2.3.). The mean age was 21.8 (SD: 0.47), and the sex ratio was 74 men to 70 women.

### 2.2. Procedures

Participants were invited to the laboratory for blood sampling. Protocols were approved by the Zurich Cantonal Ethics Committee (KEK, BASEC No. 2017-02083), and informed consent was obtained. Trained phlebotomists from the Laboratory for Social and Neural Systems Research (SNS Lab) at the University of Zurich collected 32.5 mL of peripheral blood from the antecubital vein. Blood was stored in serum, EDTA and PAXgene RNA tubes. The serum and EDTA samples were immediately centrifuged at 2000 xg 10 min to extract serum and plasma, and all samples including PAXgene tubes were stored at −80 °C until analysis.

A total of 153 samples with sufficient mRNA/DNA were processed at LIFE & BRAIN GmbH, Bonn, Germany, for high-throughput bulk analysis. mRNA was extracted from PAXgene following the protocol previously described (Becker et al., 2022), converted to cDNA using the Lexogen QuantSeq 3^′^ mRNA FWD kit (Corley et al., 2019), and sequenced at 15M reads (1×100bp) on a NovaSeq 6000. Adapter sequences and low-quality bases were trimmed with cutadapt v1.16, and mRNA-seq data were mapped to the GRCh38 human reference genome using STAR v2.7.6a, and Salmon (Jagannath et al., 2020). Gene expression was quantified using featureCounts v2.0.1 (subread package), with quality control (QC) metrics assessed via MultiQC v1.9. The resulting mRNA gene count matrix was generated using the nf-core/rnaseq pipeline. Potential batch effects were assessed via principal component analysis, with the first four components explaining 13.15% of the variance, indicating no systematic clustering (Leek et al., 2010).

DNA was isolated from EDTA blood following the method previously described (Jagannath et al., 2020). Isolated DNA was genotyped using the Illumina Infinium GSA Array mapped to GRCh38. PED files were subjected to basic QC (maf > 0.01; info > 0.8), strand flipping, sex check, and clumping (Choi, 2020). PRS, were calculated from summary statistics of recognized GWAS in European populations (ADHD: Demontis et al., 2023; PD: Forstner et al., 2021, 2023; GAD: Otowa et al., 2016, 2016; MDD: Wray et al., 2018) lifted to GRCh38 using the UCSC LiftOver tool. Principal components of genotyping were calculated, with both analyses conducted using PLINK 1.90b7 (Purcell et al., 2007) and GNU awk 5.1.0.

Blood plasma cytokines were quantified in triplicates using the Bio-Plex Human Cytokine Screening Panel, 48-Plex (BioRad, #12007283), following the manufacturer’s protocols. Raw fluorescence data were processed in Bio-Plex Manager version 6.1 (Bio-Rad Laboratories, 2023) to generate cytokine concentrations using standard curves, and subsequent analyses were conducted in R. The experimenter and wet lab personnel were blinded to PV conditions, with only the data analyst informed of group assignments.

### 2.3. Peer victimization profile

From waves four (age 11) to eight (age 20) of the z-proso panel, a binary treatment variable was constructed using self-reported data on social peer adversity, including exclusion, insult, physical violence, and property damage. Items were measured on a Likert scale to assess incidence over the last 12 months (1 = never, 2 = 1 time, 3 = 3 times, 4 = about once a month, 5 = about once a week, 6 = about every day). In each wave, the same questions had been asked regarding how much adversity participants had experienced (subordinate experiences, SE) and how much adversity they had exerted on others (dominant behavior, DB). Composite scores for SE and DB were created by summing up the respective Likert-scale variables for each adversity type in each wave. Next, totals were normalized to derive scores quantifying social peer adversity. To classify individuals as severely affected from SE or DB, the 90th percentile and above for each survey year was used (SE: 2009 = 0.36, 2011 = 0.36, 2013 = 0.28, 2015 = 0.20, 2018 = 0.20; DB: 2009 = 0.24, 2011 = 0.36, 2013 = 0.28, 2015 = 0.20, 2018 = 0.16) to remove trends and dynamically capture individuals with the most severe experiences with presumably significant impacts on their well-being. Ultimately, 16 individuals (11.1% of the analytic sample) were identified as being exclusively exposed to SE without exhibiting DB, and a binary variable was created to represent this PV profile. Instead of directly assessing the extent or proximity of PV, these individuals were thus assumed to have consistently experienced ‘pure’ victimization, even in years without assessments (Fig. S1A visualizes the derivation of the PV profile). Table 1 presents initial differences between the PV and control groups, including absolute standardized differences pre- and post-IPW.

**Table 1.** Baseline characteristics of controls and PV victims, with adjusted standardized differences (ASD).

| Variable | Control |  | PV |  | *p* | ASD<sub>0</sub> | ASD<sub>1</sub> |
| --- | --- | --- | --- | --- | --- | --- | --- |
|  | Mean | SD | Mean | SD |  |  |  |
| PV baseline* | 0.457 | 0.50 | 0.562 | 0.51 | 0.5970 | 0.497 | 0.013 |
| Corporal punishment | 0.374 | 0.41 | 0.569 | 0.58 | 0.2060 | 0.442 | 0.137 |
| Neg. behavior index | 0.282 | 0.11 | 0.235 | 0.08 | 0.0550 | 0.380 | 0.031 |
| Immigration* | 0.411 | 0.49 | 0.625 | 0.50 | 0.1160 | 0.339 | 0.040 |
| Conflictual families* | 2.225 | 1.01 | 2.062 | 1.12 | 0.5700 | 0.330 | 0.113 |
| Female* | 0.481 | 0.50 | 0.500 | 0.52 | 1.0000 | 0.305 | 0.005 |
| Genetic Variation PC2 | -0.002 | 0.08 | 0.024 | 0.05 | 0.0890 | 0.288 | 0.009 |
| Aggression score | 0.146 | 0.63 | -0.016 | 0.57 | 0.3040 | 0.261 | 0.063 |
| ADHD PRS | 0.001 | <0.01 | 0.001 | <0.01 | 0.7930 | 0.260 | 0.023 |
| Conflict resolution* | 0.519 | 0.50 | 0.562 | 0.51 | 0.7950 | 0.249 | 0.001 |
| Genetic Variation PC3 | 0.002 | 0.08 | -0.029 | 0.05 | 0.0550 | 0.202 | 0.054 |
| Healthy leisure activities | 0.601 | 0.14 | 0.578 | 0.11 | 0.4400 | 0.181 | 0.133 |
| Birth complications* | 0.194 | 0.40 | 0.188 | 0.40 | 1.0000 | 0.176 | 0.026 |
| Broken family* | 0.225 | 0.42 | 0.188 | 0.40 | 0.7700 | 0.159 | 0.022 |
| Panic disorder PRS | -0.004 | <0.01 | -0.005 | <0.01 | 0.5430 | 0.146 | 0.014 |
| Number of siblings* | 1.109 | 0.89 | 1.062 | 1.00 | 0.1670 | 0.136 | 0.054 |
| SES | 0.512 | 0.19 | 0.512 | 0.20 | 0.9960 | 0.134 | 0.071 |
| ADHD score | 0.068 | 0.73 | 0.082 | 0.49 | 0.9220 | 0.104 | 0.033 |
| Hostile attribution* | 0.426 | 0.50 | 0.312 | 0.48 | 0.4330 | 0.099 | 0.003 |
| Genetic Variation PC1 | -0.006 | 0.08 | 0.001 | 0.08 | 0.7030 | 0.089 | 0.050 |
| Adverse family incidents (n)* | 0.566 | 0.50 | 0.625 | 0.50 | 0.7900 | 0.084 | 0.005 |
| Locus of control | 2.815 | 0.54 | 2.781 | 0.58 | 0.8250 | 0.081 | 0.047 |
| Teacher rated PV baseline* | 1.302 | 0.58 | 1.312 | 1.01 | 0.0840 | 0.080 | 0.036 |
| Adv. family incidents (var)* | 1.264 | 1.44 | 1.062 | 1.44 | 0.6040 | 0.079 | 0.041 |
| Isolated in class* | 0.264 | 0.44 | 0.375 | 0.50 | 0.3800 | 0.078 | 0.012 |
| Major depression PRS | -0.007 | <0.01 | -0.007 | <0.01 | 0.6820 | 0.066 | 0.077 |
| Facial emotion recognition | 0.713 | 0.10 | 0.714 | 0.10 | 0.9890 | 0.062 | 0.020 |
| Violent peers* | 0.450 | 0.50 | 0.500 | 0.52 | 0.7920 | 0.061 | 0.003 |
| Birth rank | 1.562 | 0.79 | 1.562 | 0.81 | 0.9980 | 0.057 | 0.006 |
| Anxiety disorder PRS | -0.005 | <0.01 | -0.005 | <0.01 | 0.6720 | 0.056 | 0.049 |
| Anxiety score | 0.068 | 0.59 | 0.061 | 0.63 | 0.9670 | 0.040 | 0.018 |
| Genetic Variation PC4 | -0.005 | 0.08 | 0.007 | 0.03 | 0.2380 | 0.027 | 0.030 |
| Weight at baseline | 0.089 | 0.97 | 0.096 | 1.05 | 0.9820 | 0.025 | 0.014 |
| German lang. native* | 0.558 | 0.50 | 0.500 | 0.52 | 0.7910 | 0.016 | 0.037 |
| Age | 21.807 | 0.47 | 21.737 | 0.53 | 0.6160 | 0.004 | 0.192 |
| Coparenting index | 4.377 | 0.64 | 4.598 | 0.53 | 0.1430 | 0.004 | 0.098 |

\*Ordinal variables. *Note:* This table presents the mean (SD) for each variable in the control and PV groups included in the IPW model. P values were calculated using t-tests for continuous variables and $\chi^2$ -tests for ordinal variables. Absolute standardized differences (ASD) were computed based on pooled standard deviations for continuous variables and pooled proportions for ordinal variables, shown before (ASD<sub>0</sub>) and after IPW balancing (ASD<sub>1</sub>). Although means and SDs are displayed here for consistency, the correct methods were applied to assess balance for each variable type. It is important to note that t-tests are sensitive to sample size and may not indicate statistical significance even in the presence of covariate imbalance (Austin, 2009; Ho et al., 2007). See Fig. S1B for a visualization of the ASDs.

### 2.4. Inverse probability weighting and time-varying confounders

In our analyses, IPW (Hernán and Robins, 2020) was applied to adjust for biopsychosocial confounders related to treatment (PV) and stratified sample selection. By weighting controls to balance characteristics, IPW creates a pseudo-population that approximates a randomized trial, enhancing causal inference. This method is widely recommended for observational studies to adjust for confounding, particularly in PV research (Baldwin et al., 2023) and is effective in genomic analyses for improving variance estimation reliability (Reifeis et al., 2020). Specifically, a logit model was applied to confounders associated with both the treatment and outcome (Rubin, 1997) to derive propensity scores for PV. These scores were used to calculate weights for estimating the average treatment effect of the treated (ATT, Eq. 1),

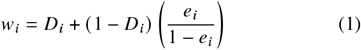

where *D* denotes the indicator of victim status and *e* is the propensity score (Morgan and Winship, 2015). The ATT was chosen to estimate the effect of PV on those directly exposed, as they represent the primary population of interest. Weights were trimmed at the 5th and 95th percentiles to reduce the influence of extreme values. Baseline data for the propensity score model were drawn from reports by the young person, teachers, and parents during the first three z-proso waves (ages 7–9), prior to measuring victimization. Missing data (0.5–6% across variables) were addressed using random forest imputation (missRanger 2.6.0; Stekhoven and Buhlmann, 2012). PRS for ADHD, GAD, MDD, and PD were included (detailed in Section 2.2.). To account for population structure within the sample, principal components from genotyping were included, with the first four selected based on eigenvalue analysis, which showed a clear inflection point at the fourth component with an eigenvalue near 1 (Fig. S3; Cattell, 1966; Price et al., 2006). See Table S2 for a complete list of the confounding factors and rationale. To minimize extrapolation bias, ATT estimation was restricted to victims with propensity scores within the range of the controls (common support) and excluded eight of the initial 24 victims with valid mRNA due to lacking suitable matches (Austin and Stuart, 2017; Stuart, 2010), yielding the final group of 16 victims. In complex survey designs (Lumley, 2023), IPW weights *w* were used as sampling weights throughout the analyses.

### 2.5. Data analysis

The aim of the analyses was to investigate the consequences of PV on biological stress indicators via (1) *in silico* cell-type deconvolution, (2) differential expression and TF analyses, and (3) cytokine dysregulation. Statistical analyses were performed using R 4.4.1 (R Development Core Team, 2023). A base model (Eq. 2) included the PV dummy with adjustments for subjects involved in DB or both SE and DB. Sexual harassment victimization (sexv), perpetration (sexp), and sexual assault victimization (assltv) were adjusted for but excluded from the initial PV profile due to distinct causal mechanisms (Frøyland et al., 2023). Adjustments were made for time-varying confounders (*X*γ) including body-mass-index (Puhl et al., 2011), exercise (Harrison and Baune, 2014; O’Connor et al., 2021; Schoeler et al., 2018), psychological help-seeking (Antoni et al., 2012; Harrison and Baune, 2014), and psychoactive substance use (Lovallo et al., 2000; O’Connor et al., 2021); to enhance statistical power, principal components were derived for each domain, reducing dimensionality while capturing about 75% of the variance. Diurnal and annual variation were controlled by including sine and cosine terms: for diurnal variation, sin (2*π*/365 × daytime) and cos (2*π*/365 × daytime); for annual variation, sin (2*π*/365 × doy) and cos (2*π*/365 × doy). All models accounted for chronic and metabolic diseases, and long-term medication use (binary). The model equation, weighted by *w* (Eq. 1), is as follows:

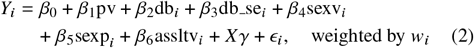

In an extended model, principal components from genes *CD14, CD19, CD3D, CD4, CD8A, FCGR3A*, and *NCAM1* were included to control for leukocyte composition, based on prior research (Cole et al., 2007; Fredrickson et al., 2013; Li et al., 2020). P values were adjusted for multiple comparisons using the false discovery rate (FDR) method (Benjamini and Hochberg, 1995) to control for type I errors.

#### 2.5.1. In silico deconvolution

Potential differences in leukocyte composition between victims and controls were assessed using an in silico cell-type deconvolution with the FARDEEP algorithm (Avila Cobos et al., 2020; Hao et al., 2019). The primary analysis used the LM22 basis matrix (Newman et al., 2015) for its more detailed cell reference profile, which surpasses that of quanTIseq. Despite being designed for microarray data, LM22 was deemed suitable due to similarities in transcript quantification between microarrays and 3’ RNA-seq, along with the latter’s broader dynamic range and enhanced detection of low-abundance transcripts (Corley et al., 2019). Results were additionally compared with quanTIseq, optimized for RNA-seq (Finotello et al., 2019), to assess agreement.

LM22 (and similarly quanTIseq) includes expression data from cells not found in peripheral blood but whose precursors are. For instance, monocytes in peripheral blood, before migrating to tissues and differentiating into M0 macrophages, exhibit precursor characteristics that influence polarization into M1 or M2 macrophages (M1, M2; Boyette et al., 2017). Consequently, M1 and M2 signals were interpreted as originating from monocytes and are referred to as M1-like and M2-like monocytes. Similarly, follicular T helper cell (Th) signals were interpreted as originating from follicular-like Th.

#### 2.5.2. Gene expression analysis

The mRNA count matrix was normalized using edgeR 4.2.2 TMM normalization (Robinson et al., 2010), with a minimum count of 5 and a proportion of 5%, followed by log2-transformation resulting in a matrix of 17,333 genes. DE genes were analyzed through two approaches: an a priori analysis focusing on CTRA genes and preselected Reactome pathways (v88; Gillespie et al., 2022) with known CTRA activity, and an exploratory analysis querying genome-wide Reactome pathway enrichment.

##### 2.5.2.1. A priori analysis

Following established methods (Li et al., 2020; Ross et al., 2021), CTRA genes (Cole, 2019) were analyzed as repeated measures, with sign-inverted counts for genes expected to be stress-downregulated. Analyses employed a complex survey design (Lumley, 2023) incorporating IPW and subject-specific intercepts. Genome-wide DE analysis was then performed using limma (Gordon K. Smyth et al., 2022) with IPW applied. Variance adjustments for the IPW pseudo-population were calculated using empirical sandwich estimators from geex v1.1.1 (Reifeis and Hudgens, 2022; Reifeis et al., 2020; Saul and Hudgens, 2020)). Significantly DE genes (|log_2_ (FC)| > log_2_ 1.2, *p* < 0.05) were analyzed via Fisher’s exact tests to identify overrepresentation in top-level Reactome pathways containing CTRA genes, including *Hemostasis* (R-HSA-109582), *Immune System* (R-HSA-1430728), *Metabolism* (R-HSA-162582), and *Signal Transduction* (R-HSA-168256), with a particular focus on antiviral and pro-inflammatory subpathways. Importantly, only 43 of the 61 CTRA genes are currently annotated in Reactome. P values were FDR-adjusted within the preselected pathways (*p* < 0.05).

##### 2.5.2.2. Exploratory analysis

Significantly up- and downregulated DE genes were used to query pathway enrichment across the entire Reactome database (annotations dated 2024-01-25) via the gprofiler2 API (Raudvere et al., 2019), to identify significant pathways across all hierarchical levels. Enrichment significance was FDR-corrected across 2,562 Reactome pathways (*p* < 0.05).

#### 2.5.3. Transcription factor activity analysis

Activity of TFs was inferred from predicted transcription factor binding sites in significantly DE genes using the TRANSFAC^®^ database (Matys et al., 2003) via gprofiler2, focusing on high-confidence matches within ±1 kb of the transcription start site, categorized as match class ±1 to ensure credibility. The Harmonizome database (Rouillard et al., 2016) was then queried to retrieve TF target genes (TFTGs). Significantly DE TFTGs were mapped to genes from Reactome-wide identified pathways to calculate the pooled gene ratio of TFs and identify potential TF–pathway associations.

#### 2.5.4. Cytokine analysis

Absolute concentrations of 16 cytokines (CCL11, CCL27, CCL4, CCL5, CLEC11A, CSF1, CXCL10, CXCL12, CXCL9, HGF, IL16, IL9, KITLG, LTA, PDGFB, and TNF) were obtained from multiplex immunoassays. Cytokine data were log2-transformed and analyzed using IPW-adjusted regression models. Statistical significance was determined with FDR-corrected p values (*p* < 0.05).

## 3. Results

### 3.1. IPW and model performance

The maximum absolute standardized difference between treated and controls for IPW covariates was 0.194, indicating satisfactory balance for the sample size (Austin, 2009). The logistic regression model generated valid propensity scores without perfect predictions. Fig. S1B shows improved balance after weighting, and Fig. S3 illustrates sufficient overlap in propensity scores between groups. Variance inflation was assessed, showing no evidence of multicollinearity or inflated standard errors. The difference between the mean of the estimated weights and their expected value was −0.009, reflecting high agreement and stability of the weighting procedure (Reifeis and Hudgens, 2022).

### 3.2. In silico deconvolution

*In silico* cell-type deconvolution revealed several differences in the leukocyte profile of victims compared to controls, suggesting the inclusion of leukocyte-encoding genes in subsequent analyses. The following coefficients reflect the relative abundance of the respective cell type. Deconvolution estimated lower proportions of M1-like monocytes (log2FC = 0.027; 95% CI = [−0.040, −0.014]; *p* = 0.0008), eosinophils (log2FC = −0.025; 95% CI = [−0.035, −0.015]; *p* = 0.0001), and follicular-like Th cells (log2FC = −0.043; 95% CI = [−0.071, −0.015]; *p* = 0.0221) in victims (Fig. 1A). The model also estimated higher proportions of activated dendritic cells (DC^+^) (log2FC = 0.036; 95% CI = [0.002, 0.071]; *p* = 0.095), although this difference did not reach statistical significance. LM22-derived estimates for M1-like monocytes and neutrophils showed similar significance patterns to supplementary quanTIseq results, supporting robustness (Table S3, Table S4).

**Figure 1:**
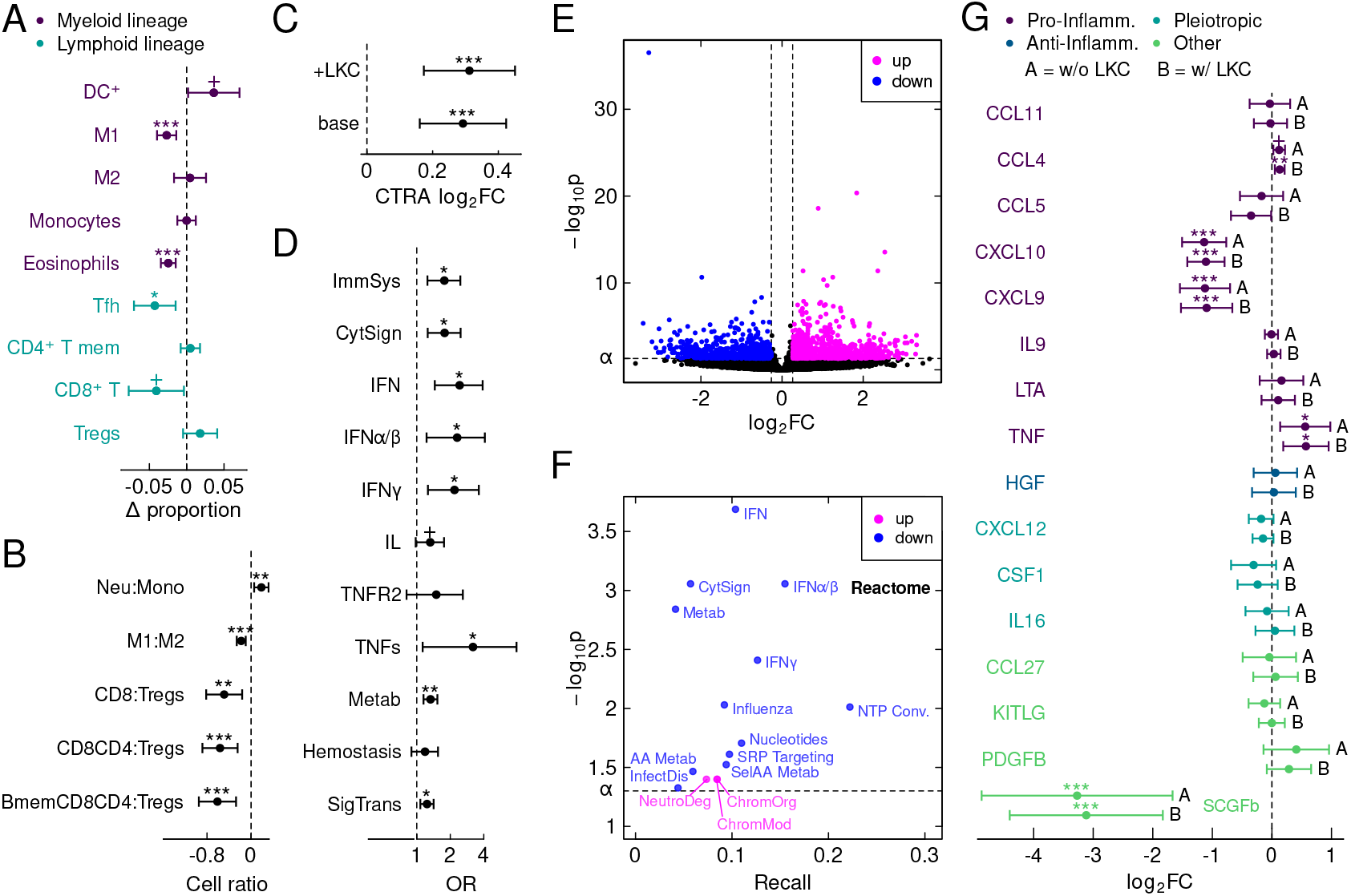
Leukocytes and gene expression. ***p < 0.001, **p < 0.01, *p < 0.05, ^+^p < 0.1. ***(A)*** Changes in relative leukocyte proportions in victims compared to controls with 95% CIs. DC^+^ = activated dendritic cells, M1 and M2 = M1-like and M2-like monocytes, Tfh = follicular-like Th, CD4^+^ T mem = memory CD4^+^ T cells, CD8^+^ T = CD8^+^ T cells, Tregs = regulatory T cells. ***(B)*** Ratios of specific cell types in the deconvolution analysis with 95% CIs. Neu = neutrophils, mono=monocytes, Bmem = memory B cells. ***(C)*** Upregulation of the CTRA profile compared to controls with 95% CIs. base=base model, +LKC=leukocyte encoding genes added in model. ***(D)*** Odds ratios for enrichment in preselected Reactome pathways from Fisher’s exact tests with 95% CIs. The x-axis is on a log scale. ***(E)*** Volcano plot of DE genes from genome-wide analysis. ***(F)*** Pathways identified from the Reactome-wide query. ***(G)*** Comparison of absolute cytokine levels in victims vs. controls with 95% CIs. A=base model, B=leukocyte encoding genes included in model. ChromMod = *Chromatin modifying enzymes*, ChromOrg = *Chromatin organization*, CytSign = *Cytokine Signaling in Immune system*, ImmSys = *Immune System*, InfectDis = *Infectious disease*, Influenza = *Influenza Infection*, InnateImm = *Innate Immune System*, NTP Conv. = *Interconversion of nucleotide di- and triphosphates*, IFNα/β = *Interferon alpha/beta signaling*, IFNγ = *Interferon gamma signaling*, IFN = *Interferon Signaling*, Metab = *Metabolism*, AA Metab = *Metabolism of amino acids and derivatives*, Nucleotides = *Metabolism of nucleotides*, Protein Metab = *Metabolism of proteins*, NeutroDeg = *Neutrophil degranulation*, SelAA Metab = *Selenoamino acid metabolism*, SigTrans = *Signal Transduction*, IL = *Signaling by Interleukins*, SRP Targeting = *SRP-dependent cotranslational protein targeting to membrane*, TNFR2 = *TNFR2 non-canonical NF-kB pathway*, TNFs = *TNFs bind their physiological receptors*, Translat = *Translation*, ViralPath = *Viral Infection Pathways*.

Several ratios of certain cell types differed in victims compared to controls. A higher ratio was observed for neutrophils:monocytes (log2FC = 0.187; 95% CI = [0.558, 0.318]; *p* = 0.0056), while lower ratios were observed for M1:M2-like monocytes (log2FC = −0.179; 95% CI = [−0.260, −0.098]; *p* = 0.0001), CD8^+^ T cells:Tregs (log2FC = −0.489; 95% CI = [−0.815, −0.162]; *p* = 0.0046), (CD8^+^ T, CD4^+^ T cells):Tregs (log2FC = −0.564; 95% CI = [−0.833, −0.244]; *p* = 0.0011), and (B memory, CD8^+^ T, CD4^+^ T cells):Tregs (log2FC = −0.611; 95% CI = [−0.949, −0.273]; *p* = 0.0011) (Fig. 1B).

The resulting M2-like skew might contribute to conditions such as autoimmune reactions and various allergic and skin disorders (Martinez et al., 2009). Indeed, post hoc analyses of average marginal effects derived from IPW-weighted quasibinomial logistic regression indicated that victims had an 18.6% higher probability of reporting allergies, including asthma, compared to controls (AME = 0.186; 95% CI = [0.188, 0.543]; *p* = 0.0405). We also noted frequent reports of skin problems among victims (results not shown).

### 3.3. Differential gene expression

The CTRA profile of victims was upregulated compared to controls, with victims in the base model showing an average log2FC of 0.292 (95% CI = [0.161, 0.424], *p* < 0.0001), equivalent to a fold change of approximately 1.22 (Fig. 1B). After controlling for leukocyte-encoding genes, the extended model indicated a slightly higher log2FC of 0.312 (95% CI = [0.173, 0.451]; *p* < 0.0001), corresponding to a fold change of about 1.24. In the genome-wide DE analysis, victims exhibited 817 up-regulated and 591 downregulated genes (|log_2_ (FC) | > log_2_ 1.2 and *p* < 0.05) (Fig. 1E). This analysis confirmed repeated measures with 26 of 61 (42.6%) DE CTRA genes in victims. Among these, 4 of 25 (16.0%) typically stress-upregulated genes were up-regulated, 18 of 36 (50.0%) typically stress-downregulated genes were downregulated, and four did not follow the expected pattern (Fig. S4). This profile suggests a skew in victims toward the typical stress-downregulated antiviral component of the CTRA.

Next, odds ratios (ORs) of Fisher’s exact tests revealed a significant overrepresentation of DE genes in several preselected Reactome pathways among victims, compared to controls. Notably, two prominent subpathways were identified within the *Immune system*: (1) one encompassing *Cytokine signaling in Immune system* (OR = 1.768; 95% CI = [1.252, 1.475]; *p* = 0.0110) and *Interferon Signaling* (OR = 2.427; 95% CI = [1.451, 3.907]; *p* = 0.0110), and culminating in *Interferon alpha/beta signaling* (OR = 2.305; 95% CI = [1.223, 4.090]; *p* = 0.0243) and *Interferon gamma signaling* (OR = 2.182; 95% CI = [1.259, 3.611]; *p* = 0.0188) activity. *Interferon alpha/beta signaling* included 11 DE genes, all downregulated, such as *IFI27* (log2FC = −2.142; *p* = 0.0010). *Interferon gamma signaling* included 14 DE genes: one upregulated (*ICAM1*, log2FC = 0.516; *p* = 0.0243) and 13 downregulated (e.g., *TRIM10*; log2FC = −1.098; *p* = 0.0233). (2) The other subpathway culminated in *TNFs bind their physiological receptors* (OR = 3.199; 95% CI = [1.128, 7.883]; *p* = 0.0333), which included 5 DE genes: two upregulated (e.g., *TNFRSF9*; log2FC = 1.327, *p* = 0.0049) and three downregulated (e.g., *FASLG*; log2FC = −2.324, *p* = 0.0008). Two top-level pathways also showed significant overrepresentation of DE genes: *Metabolism* (OR = 1.328; 95% CI = [1.148, 1.531]; *p* = 0.0077) and *Signal transduction* (OR = 1.236; 95% CI = [1.076, 1.416]; *p* = 0.0135). *Metabolism* included 174 DE genes, 88 upregulated (e.g., *TPH1*; log2FC = 2.652, *p* = 0.0010) and 86 downregulated (e.g., *FHL2*; log2FC = 2.916, *p* = 0.0073). *Signal transduction* included 191 DE genes, 123 upregulated (e.g., *FPR3*; log2FC = 3.314, *p* = 0.0001), and 68 downregulated (e.g., *H2BC7*; log2FC = −2.780, *p* < 0.0001). While *Signaling by Interleukins* did not reach statistical significance (OR = 1.320; 95% CI = [0.977, 1.759]; *p* = 0.0789), the pathway included 40 DE genes, 19 upregulated (e.g., *S100B*; log2FC = 2.865, *p* = 0.0459) and 21 downregulated (e.g., *FASLG*). (Fig. 1D) presents the ORs summarized in Table 2, with the *a priori* pathways included in the combined pathway hierarchy illustrated in (Fig. 2).

**Table 2.** Overrepresentation analysis of Reactome pathways based on odds ratios (OR) for differentially expressed genes.

| Pathway | OR | Lower | Upper | $p_{adj.}$ |
| --- | --- | --- | --- | --- |
| Immune System | 1.768 | 1.246 | 2.462 | 0.0110 |
| Cytokine signaling in Immune System (R-HSA-1280215) | 1.778 | 1.252 | 2.475 | 0.0110 |
| Interferon signaling (R-HSA-913531) | 2.427 | 1.451 | 3.907 | 0.0110 |
| Interferon alpha/beta signaling (R-HSA-909733) | 2.305 | 1.223 | 4.090 | 0.0243 |
| Interferon gamma signaling (R-HSA-877300) | 2.181 | 1.259 | 3.611 | 0.0188 |
| Signaling by interleukins (R-HSA-449147) | 1.320 | 0.977 | 1.757 | 0.0789 |
| TNFR2 non-canonical NF-κB pathway (R-HSA-5668541) | 1.496 | 0.808 | 2.594 | 0.1578 |
| TNFs bind their physiological receptors (R-HSA-5669034) | 3.199 | 1.128 | 7.883 | 0.0458 |
| Metabolism | 1.328 | 1.148 | 1.531 | 0.0077 |
| Hemostasis | 1.185 | 0.894 | 1.550 | 0.1626 |
| Signal Transduction | 1.236 | 1.075 | 1.416 | 0.0135 |

The table lists pathways with their odds ratios (OR), lower and upper confidence intervals, and adjusted $p$ values ( $p_{adj.}$ ). Indentation reflects the hierarchical structure of Reactome pathways.

**Figure 2:**
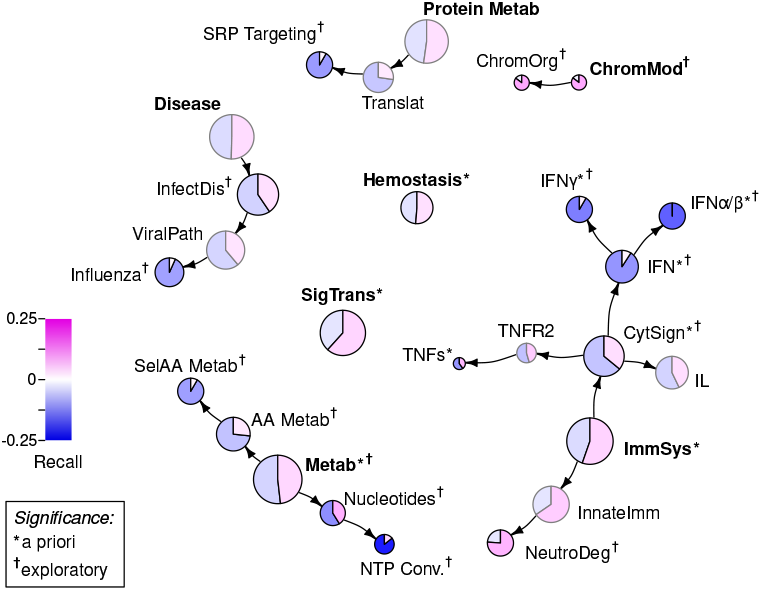
Hierarchy of queried Reactome pathways. ^*,†^ *p* < 0.05. The figure displays the hierarchical organization of pathways with overrepresentation in up- and downregulated genes, as visualized in the Reactome pathway browser. Node sizes correspond to relative term sizes, and colors indicate the gene ratio (‘recall’ R), representing the proportion of DE genes in a pathway that are present in the query set. Magenta shades indicate upregulation, while blue shades denote downregulation. The figure integrates results from both *a priori* and exploratory Reactome analyses. Abbreviations are detailed in the caption of Fig. 1.

The exploratory Reactome-wide query confirmed and extended *a priori* findings. For each pathway, the gene-set ratio (also referred to as ‘recall’, *R*), representing the proportion of DE genes within the pathway present in the query set, along with FDR-adjusted p values, were derived. This analysis identified an overrepresentation of upregulated genes in pathways associated with *Chromatin organization* (R-HSA-4839726) and *Immune System* (R-HSA-168256), while downregulated genes were over-represented in *Metabolism* (R-HSA-1430728), *Immune System* (R-HSA-168256), *Disease* (R-HSA-1643685), and *Metabolism of proteins* (R-HSA-392499) in victims, compared to controls (Table 3; Fig. 1F). Top-level *Immune System* was not significant in the Reactome-wide query; however, its significance in the *a priori* tests suggests statistical dilution due to its large gene set. These results indicated that the *a priori* cytokine- and interferon-related pathways were downregulated. In contrast, another pathway within the *Immune System, Neutrophil degranulation* (R-HSA-6798695), was upregulated. Two new pathways were identified within *Metabolism*: one culminating in *Interconversion of Nucleotide Di- and Triphosphates* (R-HSA-499943) and another in *Selenoamino acid metabolism* (R-HSA-2408522), both of which showed an overrepresentation of downregulated genes. Additionally, the top-level pathway *Chromatin Organization* (R-HSA-4839726) and its subpathway *Chromatin Modifying Enzymes* (R-HSA-3247509) exhibited an overrepresentation of upregulated genes. Although the top-level pathways *Metabolism of proteins* and *Disease* were not significant, *SRP-dependent cotranslational protein targeting to membrane* (R-HSA-1799339) and *Influenza Infection* (R-HSA-168256) demonstrated significant downregulation, suggesting that statistical dilution of large gene sets may have obscured significance at the top level.

**Table 3.**
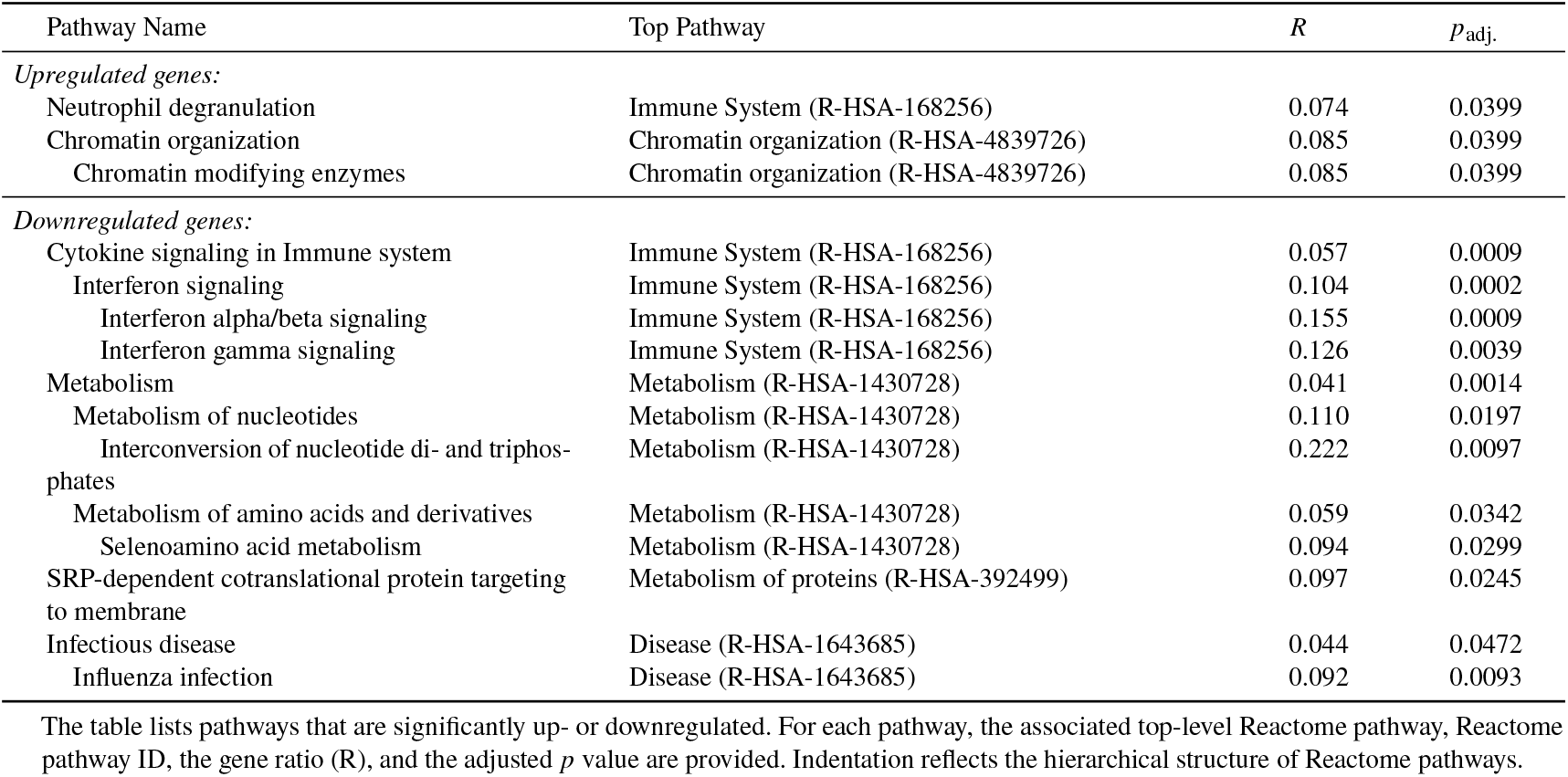
Overrepresentation analysis of *exploratory* Reactome pathways for differentially expressed genes.

Among these findings, two key components of interferon-related signaling showed significance in the DE gene analysis, including *STAT1* (downregulated, log2FC = −0.486, *p* = 0.0008) and *SOCS4* (upregulated, log2FC =1.223, *p* = 0.0026), while *JAK1* was not significant (log2FC = 0.125, *p* = 0.2790). Furthermore, the glucocorticoid receptor gene *NR3C1* (log2FC = 0.280, *p* = 0.0414) and *NFKB1*, which encodes NF-κB (log2FC = 0.214, *p* = 0.0230), were upregulated. Fig. 2 presents a targeted excerpt from the Reactome browser, highlighting key pathways and sub-pathways and their placement within the broader Reactome hierarchy. Notably, *a priori* pathways and sub-pathways within *Immune System* and *Metabolism*—where CTRA genes are known to be active—were also identified in the exploratory Reactome query after correction for multiple testing across more than 2,500 pathways.

### 3.4. Transcription factors

A total of 108 active TFs were inferred, with 93 associated with upregulated DE genes and 15 with downregulated DE genes in victims, compared to controls (Fig. 3A). Recall (R), defined here as the proportion of DE genes with TF binding sites matching their respective TFs, was used to evaluate motif overlap. To identify a subset of the most relevant TFs, a mean plus one standard deviation was applied as a reference threshold, selecting three TFs associated with downregulated genes: IRF2 (R = 0.088; *p* = 0.0144), IRF8 (R = 0.114; *p* = 0.0028), and STAT2 (R = 0.364; *p* = 0.0166). To avoid overlooking potentially meaningful signals, two additional TFs below the threshold were included: IRF4 (R = 0.055; *p* = 0.0144), associated with upregulated genes and included due to its membership in the IRF family, and NF-κB (R = 0.040; *p* = 0.0435), associated with downregulated genes and included because of its prominent role in inflammation. All five TFs had matches with TFTGs in Reactome pathways of the exploratory analysis, with STAT2 and IRF2 regulating the largest number of targets (Fig. 3B). Upregulated Chromatin organization and downregulated Interferon signaling pathways emerged as most strongly regulated by the selected TFs. No substantial evidence indicated that TF activity was regulated by the expression of their encoding genes (Fig. 3C). This conclusion is supported by non-significant p values for *IRF2* (log2FC = 0.071; *p* = 0.7232), *IRF4* (log2FC = −0.799; *p* = 0.5141), *IRF8* (log2FC = −0.818; *p* = 0.2740), and *STAT2* (log2FC = 0.959; *p* = 0.8330). Although *NFKB1* showed a significant p value (log2FC = 1.160; *p* = 0.0230), the fold change was minimal. Notably, *NFKB1* was the only DE TFTG within the CTRA gene set.

**Figure 3:**
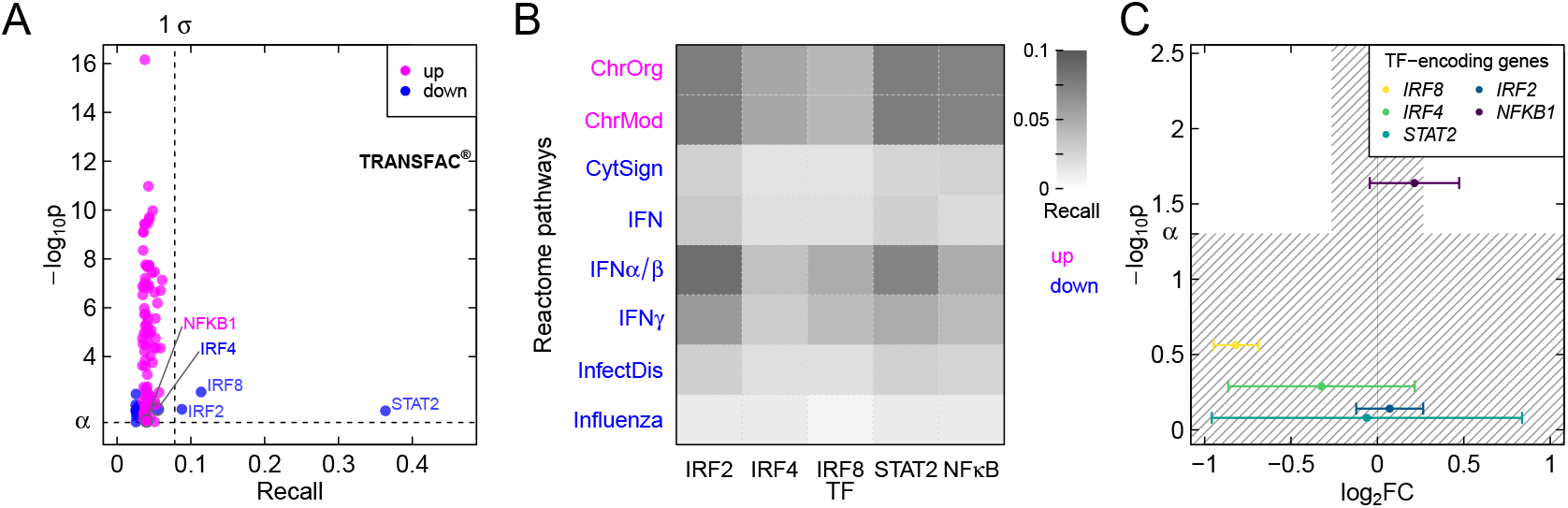
Transcription factors (TFs). ***(A)*** TF yield of TRANSFAC^®^ query, up = TFs associated with upregulated DE genes, down = TFs associated with downregulated DE genes. The vertical dashed line denoting 1 SD above the mean gene ratio (R), defined as the proportion of DE genes with TF binding sites matching their respective TFs. ***(B)*** TFs with TFTG matches in Reactome query pathways. Only pathways with reasonable matches (R > 0.01) are shown. Gray intensity indicates gene ratio (‘recall’ R) within the pathways, defined as the proportion of DE genes in a pathway relative to its total genes, with magenta representing upregulated and blue representing downregulated pathways. ***(C)*** Coefficient plot of TF-encoding genes, highlighting limited correspondence between TF activity and differential expression of their encoding genes, suggesting alternative regulatory mechanisms beyond transcriptional control.

### 3.5. Cytokines

In both the base model and the extended model controlling for leukocytes, cytokine levels differed in victims from those in controls, showing both elevations and reductions. While the overall trend between models remained consistent, significance increased slightly for several cytokines in the extended model. Among pro-inflammatory cytokines, elevations in the extended model included TNF (log2FC = 0.571; 95% CI = [0.186, 0.957]; *p* = 0.0130) and CCL4 (log2FC = 0.132; 95% CI = [0.049, 0.214]; *p* = 0.0087), while reductions were observed for CXCL10 (log2FC = −1.108; 95% CI = [−1.424, −0.791]; *p* < 0.0001) and CXCL9 (log2FC = −1.096; 95% CI = [−1.532, −0.659]; *p* < 0.0001). CCL5 was also reduced but did not reach statistical significance (log2FC = −0.350; 95% CI = [−0.692, −0.009]; *p* = 0.1185). Similarly, CSF1 was reduced without statistical significance (log2FC = −0.240; 95% CI = [−0.581, −0.101]; *p* = 0.2941). Among growth factors, CLEC11A was significantly reduced (log2FC = −3.116; 95% CI = [−4.416, −1.816]; *p* < 0.0001) while PDGFB showed a tendency to be elevated (log2FC = 0.286; 95% CI = [−0.090, 0.663]; *p* < 0.2687) but was not statistically significant. The mean log2FC for pro-inflammatory cytokines in the extended model was 0.217 (SD = 0.601). The average absolute change between the base and extended models in log2FC was 0.070 (SD = 0.057), while the change in BIC showed considerable variability (mean = 10.366; SD = 10.842). Fig. 1G visualizes the cytokine results, with complete cytokine level estimates provided in Table S5 and Table S6.

## 4. Discussion

PV is a common problem with documented, potentially serious consequences for mental and physical health (Lereya et al., 2015). Additionally, PV induces significant social stress during critical periods of immune system development (Rowson et al., 2019), potentially affecting leukocyte composition, gene expression, TF activity, and cytokine levels. Leveraging longitudinal data, this study examined the biological consequences of PV in young adulthood. PV profiles were carefully derived from self-reports, focusing on ‘pure’ victims—individuals who experienced victimization but did not perpetrate it. Reports from the young person, parents, and teachers, along with genomic data, were used to implement a counterfactual design with IPW accounting for baseline and time-varying confounders across all analyses.

Adolescent PV was associated with shifts in leukocyte composition, characterized by reduced M1-like monocytes, a lower M1-like:M2-like monocyte ratio, and a trend toward increased DC^+^ proportions, consistent with prior research noting that dendritic cells and monocytes are transcriptionally sensitive to the social environment (Cole et al., 2011). The implicated skew toward an M2 phenotype reflects suppressed pro-inflammatory responses, aligning with downregulated *Interferon alpha/beta* (IFN-α/β) *signaling* and *Interferon gamma* (IFN-γ) *signaling* pathways (Mosser and Edwards, 2008; Sica and Mantovani, 2012). Note that this pattern was observed in monocytic precursors, suggesting that actual macrophages likely reside in tissues beyond the scope of peripheral blood measurements. While M2 contribute to tissue repair and anti-inflammatory processes, they also elevate autoimmune risks (Martinez et al., 2009), which aligns with the increased asthma, allergy, and skin problems in victims. Additionally, M2-like skewing suggests a transition from an acute to a chronic immunoregulatory state (Sica and Mantovani, 2012).

PV-induced immune dysregulation was further evidenced by an elevated neutrophil:monocyte ratio and upregulated NR3C1, suggesting altered glucocorticoid sensitivity as previously reported (Cole, 2008). Reduced CD8+ T cells:Tregs ratios, reflecting Treg-mediated inhibition of cytotoxic CD8+ T cells, suggests immune suppression and potential exhaustion (Haruna et al., 2022; Schietinger and Greenberg, 2014). Reduced eosinophils and follicular-like Th levels, potentially reflecting a tendency toward lower CCL5, a known eosinophil chemoattractant (Kameyoshi et al., 1992), further suggest an immunosuppressive profile and autoimmune predisposition. These findings align with stress-induced leukocyte modulation models (Cole, 2008; Dhabhar, 2009; Munshi et al., 2020).

The upregulation of the CTRA profile in victims is consistent with stress-induced observations in other social adversities, including low socioeconomic status (Knight et al., 2016; Levine et al., 2017; Powell et al., 2013; Shanahan et al., 2022), low social connection and loneliness (Chun et al., 2017; Cole et al., 2015a, 2015b; Kim et al., 2021; Korytář et al., 2016; S.-H. Lee et al., 2023; Moieni et al., 2015; Snodgrass et al., 2022, 2018, 2019), psychological stress and anxiety (Chiang et al., 2019; Wingo and Gibson, 2015), and racial and sexual discrimination (M. J. Lee et al., 2021; Li et al., 2020; Thames et al., 2019).

Reactome analyses highlighted immune, metabolic, and chromatin organization pathways in victims, suggesting potential long-term effects on immune function and physical health. *Immune System* pathways included downregulated IFN-α/β and IFN-γ signaling, consistent with reduced antiviral responses and the observed M2-like skew (Mosser and Edwards, 2008; Sica and Mantovani, 2012). Downregulated *Metabolism* pathways suggested mitochondrial dysfunction (Blanco and Kaplan, 2023; Hirschenson et al., 2022), a “hallmark of aging” (López-Otín et al., 2013), potentially linked to cellular energy deficits, fatigue, and metabolic symptoms reported by victims (Gini and Pozzoli, 2009; Hager and Leadbeater, 2016; Hirschenson et al., 2022), as well as increased type-2 diabetes risk (Haythorne et al., 2019; Prashanth et al., 2021). Upregulated *Chromatin organization* pathways, associated with transcriptional regulation changes and epigenetic DNA modifications, may amplify stress responsiveness and hint at potential cancer risks (MacAlpine and Almouzni, 2013; Zhao et al., 2021).

PV was also associated with disruptions in *Metabolism of proteins*, with downregulated *SRP-dependent cotranslational protein targeting to membrane* suggesting translational read-through and misfolded protein accumulation (Karijolich and Yu, 2014), potentially leading to cellular stress, dysfunction, and death, and contributing to inflammation and neurodegenerative diseases such as dementia, Alzheimer’s, Parkinson’s, and Huntington’s (Ciechanover and Kwon, 2015; Karanth et al., 2020).

Downregulation in *Disease* pathways, particularly *Infectious disease* and *Influenza infection*, further supports immune dysregulation resembling expression profiles observed during these conditions; NF-κB, essential for cytokine production and pathogen clearance, is often hijacked by viruses like influenza to suppress host immunity (Gaur et al., 2011). Dysregulation within enriched *Hemostasis* pathways may amplify inflammation and increase risks of thrombosis or clotting complications (Margetic, 2012), while impaired *Signal Transduction* pathways, as supported by downregulated *STAT1* and non-significant changes in *JAK1*, may contribute to immune dysregulation, autoimmunity, and cancer risk (Liongue et al., 2023; Samra et al., 2024). Together, these findings suggest that PV-induced stress may contribute to disruptions across immune, metabolic, and regulatory pathways.

Key TFs, including STAT1, IRF2, and NF-κB, exhibited activity differences in victims compared to controls, with regulation independent of encoding gene expression, suggesting roles for miRNA activity, protein-protein interactions, or post-transcriptional modifications (Arimoto et al., 2018). STAT2 and IRF2 emerged as key Interferon signaling regulators, with STAT2 likely inhibiting IFN-γ signaling and IRF2 likely inhibiting IFN-α/β signaling (Arimoto et al., 2018; Hida et al., 2000; Negishi et al., 2018; Oshima et al., 2004). The suppression of IFN-γ signaling can further promote the M2 skew with its autoimmune implications (Elser et al., 2002). STAT2 and IRF2 were also implicated as potential promoters in the upregulated *Chromatin organization* pathways, suggesting a role in driving transcriptional changes and possibly facilitating epigenetic DNA modifications. Elevated NF-κB activity, suggestive of increased CCL4 and TNF levels, likely amplifies pro-inflammatory responses under PV-related stress (Liu et al., 2017; Miller et al., 2009). By contrast, IRF8 and IRF4 appeared to play minimal roles in the identified pathways, with their regulatory activity restricted to a subset of TFTGs.

Altered plasma cytokine levels in victims compared to controls corroborated the observed immune dysregulation, with elevated TNF and CCL4 linked to PV-associated symptoms such as fatigue, anxiety, depressive symptoms, and metabolic syndrome (Munshi et al., 2020; Slavich and Irwin, 2014). In contrast, reduced CXCL10 and CXCL9 levels reflected diminished IFN-γ signaling and impaired M1 activity (Mosser and Edwards, 2008; Samra et al., 2024). Reduced CLEC11A, important for hematopoietic progenitor cell development (Hiraoka et al., 1997; Ito et al., 2003) and β-cell function (Shi et al., 2023), may exacerbate immune dysregulation and aligns with risks of type-2 diabetes linked to mitochondrial dysfunction. Finally, an elevated tendency of PDGFB, vital for tissue repair, cell proliferation, and vascular development (Vanlandewijck et al., 2015), may signify an adaptive response to stress-induced tissue damage.

Changes in leukocyte composition likely contributed to altered cytokine levels, as, for instance, macrophages and dendritic cells are primary producers of pro-inflammatory cytokines (Yoshimura et al., 2021), although reverse causal mechanisms must also be considered. Persistent cytokine alterations, evident after leukocyte composition adjustments, may suggest additional mechanisms. For instance, the downregulated pathway T*NFs bind their physiological receptors* implies impaired receptor-ligand interactions, potentially disrupting inflammatory signaling. Alternatively, *STAT1* downregulation alongside unchanged *JAK1* expression suggest disrupted negative cytokine regulation mediated by the JAK1–STAT1 pathway (Yasukawa et al., 2000), associated with autoimmune and inflammatory diseases, interferon resistance, and impaired cytokine balance, potentially shaped by prolonged stress exposure. Together, these findings highlight a PV-induced cytokine imbalance, contributing to suppressed antiviral responses, immune dysregulation, and increased risk of autoimmune conditions (Marsland et al., 2017).

Stronger evidence of suppressed antiviral functions than heightened inflammation was observed in victims compared to controls. This distinction may reflect immune tolerance or exhaustion potentially associated with the chronic stress of longstanding PV (Haruna et al., 2022; Schietinger and Greenberg, 2014). The PV profile, marked by impaired immune response, persistent dysregulation, and M2-like skew (Sica and Mantovani, 2012), as well as autoimmune tendencies, suggests a chronic PV phenotype: Initial acute phases of stress might have occurred earlier, potentially leading to an adapted or dysregulated immune state rather than an initial inflammatory response. An alternative explanation is that chronic inflammation levels may generally be low in young adulthood and so less prone than the innate antiviral response to stress-related upregulation during this developmental period.

## 5. Limitations and future work

Further investigation is needed to determine how such profiles contribute to PV-specific health risks. The detailed PV model and IPW addressed common confounding factors in PV studies. Nonetheless, the propensity score model, while addressing major confounders, may not fully account for unmeasured variables, potentially limiting causal inferences. Reverse causation—where innate traits influence both PV exposure and outcomes—remains a theoretical possibility. However, no evidence from this study supports this hypothesis. Future studies should employ larger sample sizes to enable the exploration of nuanced patterns, such as early vs. late or chronic vs. acute victimization. Moreover, incorporating longitudinal mRNA data and tissue samples beyond peripheral blood will provide deeper insights into the molecular mechanisms underlying these processes. Participants were classified as ‘victims’ using external criteria, as self-reported appraisals of social adversity were unavailable. However, appraisals are stronger predictors of biological impacts than external assessments (Kemeny, 2003; M. J. Lee et al., 2021; Zych et al., 2020b). This limitation suggests that our findings may underestimate the associations involving PV, underscoring the need for appraisal measures in future studies. The data, derived from a specialized sample in Zurich, a city with high living standards, may limit the generalizability of the findings. Nonetheless, PV predicted indicators of distress even within this privileged context, demonstrating the value of this sample for understanding the effects of social adversity. While the sample size was suitable for analyzing gene signatures and pathways, larger samples are needed to investigate smaller gene sets or individual genes in greater detail (Cole et al., 2003). Analytical tools such as WGCNA (Langfelder and Horvath, 2008) and MOFA (Pierre-Jean et al., 2020) were not used due to their lack of support for IPW; future updates should address this need.

## 6. Conclusions

This study examined the impact of a decade of PV during adolescence on gene expression, leukocyte composition, and cytokine profiles in young adulthood. Using a counterfactual design, we observed suppressed antiviral responses, heightened inflammatory markers, and a profile indicating potential autoimmune risk, consistent with prior research on social adversity. Dysregulated immune and metabolic pathways were associated with these changes and were regulated by key TFs, including STAT2, IRF2, and NF-κB. These TFs showed activity independent of their encoding genes, suggesting alternative regulatory mechanisms beyond transcriptional control. Cytokine alterations further highlighted immune dysregulation associated with PV. These findings shed light on the systemic effects of PV on immune function and its potential long-term health risks in young adulthood. Prevention and resilience-building strategies should prioritize early identification and support for at-risk adolescents, addressing both the psychological and biological consequences of PV.

## Acknowledgements

We gratefully acknowledge the voluntary participation of all study participants. During the work on this paper, Jens Heumann was a fellow of the International Max Planck Research School on the Life Course (LIFE, http://imprs-life.mpg.de).

## Ethical standards

The authors assert that all procedures contributing to this work comply with the ethical standards of the relevant national and institutional committees on human experimentation and with the Helsinki Declaration of 1975, as revised in 2008.

## Appendix

**Table S1:** List of variables and details of the 23 strata used for sample selection.

| **Stratum** | **Sex** | **Bullying** | **Aggression** | **Anxiety** | **Conflict Skills** | **n_strata** | **n_sampled** |
| --- | --- | --- | --- | --- | --- | --- | --- |
| 1 | m | FALSE | FALSE | FALSE | TRUE | 118 | 13 |
| 2 | f | FALSE | TRUE | TRUE | FALSE | 109 | 24 |
| 3 | f | FALSE | TRUE | FALSE | FALSE | 99 | 13 |
| 4 | f | FALSE | FALSE | FALSE | FALSE | 94 | 13 |
| 5 | m | FALSE | TRUE | TRUE | TRUE | 85 | 21 |
| 6 | m | FALSE | FALSE | TRUE | FALSE | 81 | 12 |
| 7 | f | FALSE | FALSE | FALSE | TRUE | 80 | 10 |
| 8 | m | FALSE | FALSE | FALSE | FALSE | 76 | 7 |
| 9 | m | FALSE | TRUE | TRUE | FALSE | 76 | 12 |
| 10 | f | FALSE | TRUE | TRUE | TRUE | 66 | 14 |
| 11 | m | FALSE | FALSE | TRUE | TRUE | 64 | 6 |
| 12 | f | FALSE | TRUE | FALSE | TRUE | 54 | 13 |
| 13 | f | FALSE | FALSE | TRUE | FALSE | 45 | 10 |
| 14 | m | FALSE | TRUE | FALSE | FALSE | 34 | 7 |
| 15 | m | FALSE | TRUE | FALSE | TRUE | 35 | 6 |
| 16 | f | FALSE | FALSE | TRUE | TRUE | 26 | 5 |
| 17 | m | TRUE | TRUE | TRUE | TRUE | 13 | 4 |
| 18 | f | TRUE | TRUE | TRUE | FALSE | 25 | 3 |
| 19 | m | TRUE | FALSE | TRUE | FALSE | 10 | 1 |
| 20 | f | TRUE | TRUE | FALSE | FALSE | 16 | 0 |
| 21 | m | TRUE | TRUE | TRUE | FALSE | 13 | 0 |
| 22 | m | TRUE | FALSE | TRUE | TRUE | 12 | 0 |
| 23 | f | TRUE | FALSE | FALSE | TRUE | 11 | 0 |

**Table S2:** Confounding factors affecting the causal pathway from PV to health-related outcomes, identified through expert knowledge. Variables include predictors of both exposure and outcome (Rubin, 1997) and incorporate interactions and higher order terms (SES^2^ and age^2^) to account for non-linear influences.

| Confounder | Source |
| --- | --- |
| ADHD-score, conflict resolution, corporal punishment, isolated in class, teacher rated bullying | K. Holt and L. Espelage, 2007; Lereya et al., 2015; Zych et al., 2020a |
| Adverse family incidents, broken family, conflictual family | Forster et al., 2020; Lereya et al., 2013 |
| Age | Eastman et al., 2018 |
| Aggression, anxiety* | Espelage et al., 2001; Schoeler et al., 2018 |
| Birth complications | DiLalla et al., 2021 |
| Birth rank | Dantchev and Wolke, 2019 |
| Coparenting<sup>‡</sup> | Martínez et al., 2019 |
| Depression* | Penner-Goeke et al., 2023; Rose et al., 2016 |
| Facial emotion recognition<sup>‡</sup> | Dede et al., 2021 |
| Genetic disposition<sup>§</sup> | Brendgen et al., 2013 |
| Healthy leisure activities<sup>¶</sup>, body weight (at baseline) | Puhl et al., 2011 |
| Hostile attribution | Perhamus et al., 2022 |
| Immigration, German language native | Qin et al., 2008 |
| Negative behavior in childhood<sup>¶</sup>, self-control | Cook et al., 2010; Jennings et al., 2011; Robson et al., 2020; Zych et al., 2020a |
| Number of siblings | Karišik, 2019; Wu et al., 2023 |
| Sex | Casper and Card, 2017; Schoeler et al., 2018; White and Kaffman, 2020 |
| Socio-economic status (SES) | Delprato et al., 2017; Shanahan et al., 2022; Tippet and Wolke, 2014; Wong and Schonlau, 2013 |
| Violent peers | Duffy and Nesdale, 2009 |

\*For anxiety and depression, PRS were used, in addition to a score for anxiety. <sup>‡</sup>Coparenting score was derived from the extent to which father and mother agreed in their responses to survey questions on parenting. <sup>‡</sup>For facial emotion recognition accuracy a score was created out of correct answers to Assessing Children's Emotional Skills (ACES; Schultz et al., 2004) instrument used in z-proso. <sup>§</sup>Healthy leisure and negative behavior scores were created from a score of corresponding z-proso questions. <sup>¶</sup>First four principal components from genotyping were used. <sup>||</sup>Socio-economic status (SES) was created from parents' income, education and job prestige (ISEI). References of other scales and instruments used in z-proso can be found in the handbook (z-proso Project Team, 2024).

**Table S3:** Leukocytes LM22.

| | $\beta$ | SE | $t$ | $p$ | $p_{adj.}$ | 2.5% | 97.5% |
| --- | --- | --- | --- | --- | --- | --- | --- |
| B.cells.memory | -0.0080 | 0.011 | -0.696 | 0.4880 | 0.5964 | -0.031 | 0.015 |
| B.cells.naive | 0.0402 | 0.015 | 2.621 | 0.0100 | 0.0522 | 0.010 | 0.071 |
| Dendritic.cells.activated | 0.0363 | 0.017 | 2.093 | 0.0387 | 0.0950 | 0.002 | 0.071 |
| Dendritic.cells.resting | 0.0000 | 0.017 | 0.002 | 0.9987 | 0.9987 | -0.033 | 0.033 |
| Eosinophils | -0.0247 | 0.005 | -4.915 | <0.0001 | 0.0001 | -0.035 | -0.015 |
| Macrophages.M0 | 0.0006 | 0.007 | 0.083 | 0.9340 | 0.9987 | -0.014 | 0.015 |
| Macrophages.M1 | -0.0268 | 0.007 | -4.120 | 0.0001 | 0.0008 | -0.040 | -0.014 |
| Macrophages.M2 | 0.0045 | 0.011 | 0.414 | 0.6799 | 0.7873 | -0.017 | 0.026 |
| Mast.cells.activated | 0.0123 | 0.013 | 0.978 | 0.3303 | 0.5590 | -0.013 | 0.037 |
| Mast.cells.resting | -0.0203 | 0.017 | -1.165 | 0.2467 | 0.4522 | -0.055 | 0.014 |
| Monocytes | -0.0002 | 0.006 | -0.025 | 0.9804 | 0.9987 | -0.013 | 0.012 |
| Neutrophils | 0.0069 | 0.009 | 0.727 | 0.4690 | 0.5964 | -0.012 | 0.026 |
| NK.cells.activated | -0.0258 | 0.012 | -2.091 | 0.0389 | 0.0950 | -0.050 | -0.001 |
| NK.cells.resting | 0.0107 | 0.015 | 0.717 | 0.4746 | 0.5964 | -0.019 | 0.040 |
| Plasma.cells | -0.0124 | 0.006 | -2.001 | 0.0478 | 0.1052 | -0.025 | 0.000 |
| T.cells.CD4.memory.activated | 0.0049 | 0.007 | 0.751 | 0.4544 | 0.5964 | -0.008 | 0.018 |
| T.cells.CD4.memory.resting | -0.0098 | 0.011 | -0.857 | 0.3934 | 0.5964 | -0.033 | 0.013 |
| T.cells.CD4.naive | 0.0467 | 0.020 | 2.311 | 0.0227 | 0.0832 | 0.007 | 0.087 |
| T.cells.CD8 | -0.0407 | 0.019 | -2.180 | 0.0314 | 0.0950 | -0.078 | -0.004 |
| T.cells.follicular.helper | -0.0427 | 0.014 | -3.035 | 0.0030 | 0.0221 | -0.071 | -0.015 |
| T.cells.gamma.delta | 0.0301 | 0.012 | 2.559 | 0.0119 | 0.0522 | 0.007 | 0.053 |
| T.cells.regulatory.Tregs. | 0.0180 | 0.012 | 1.561 | 0.1214 | 0.2428 | -0.005 | 0.041 |

Statistics for associations between leukocyte profiles and PV as shown in Fig. 1A.

**Table S4:** Leukocytes quanTIseq.

| | $\beta$ | SE | $t$ | $p$ | $p_{adj.}$ | 2.5% | 97.5% |
| --- | --- | --- | --- | --- | --- | --- | --- |
| B.cells | 0.0247 | 0.015 | 1.640 | 0.1040 | 0.1950 | -0.005 | 0.055 |
| Macrophages.M1 | -0.0315 | 0.012 | -2.580 | 0.0113 | 0.0282 | -0.056 | -0.007 |
| Macrophages.M2 | -0.0778 | 0.082 | -0.953 | 0.3430 | 0.3810 | -0.240 | 0.084 |
| Monocytes | 0.0722 | 0.025 | 2.910 | 0.0044 | 0.0147 | 0.023 | 0.121 |
| Neutrophils | 0.0759 | 0.020 | 3.740 | 0.0003 | 0.0030 | 0.036 | 0.116 |
| NK.cells | 0.0206 | 0.013 | 1.580 | 0.1170 | 0.1950 | -0.005 | 0.047 |
| T.cells.CD4 | 0.0646 | 0.047 | 1.370 | 0.1730 | 0.2340 | -0.029 | 0.158 |
| T.cells.CD8 | 0.0139 | 0.011 | 1.330 | 0.1880 | 0.2340 | -0.007 | 0.035 |
| Tregs | -0.1610 | 0.048 | -3.380 | 0.0010 | 0.0051 | -0.255 | -0.067 |
| Dendritic.cells | -0.0016 | 0.024 | -0.067 | 0.9460 | 0.9460 | -0.049 | 0.046 |

Statistics for associations between leukocyte profiles and PV using quanTIseq.

**Figure S1:**
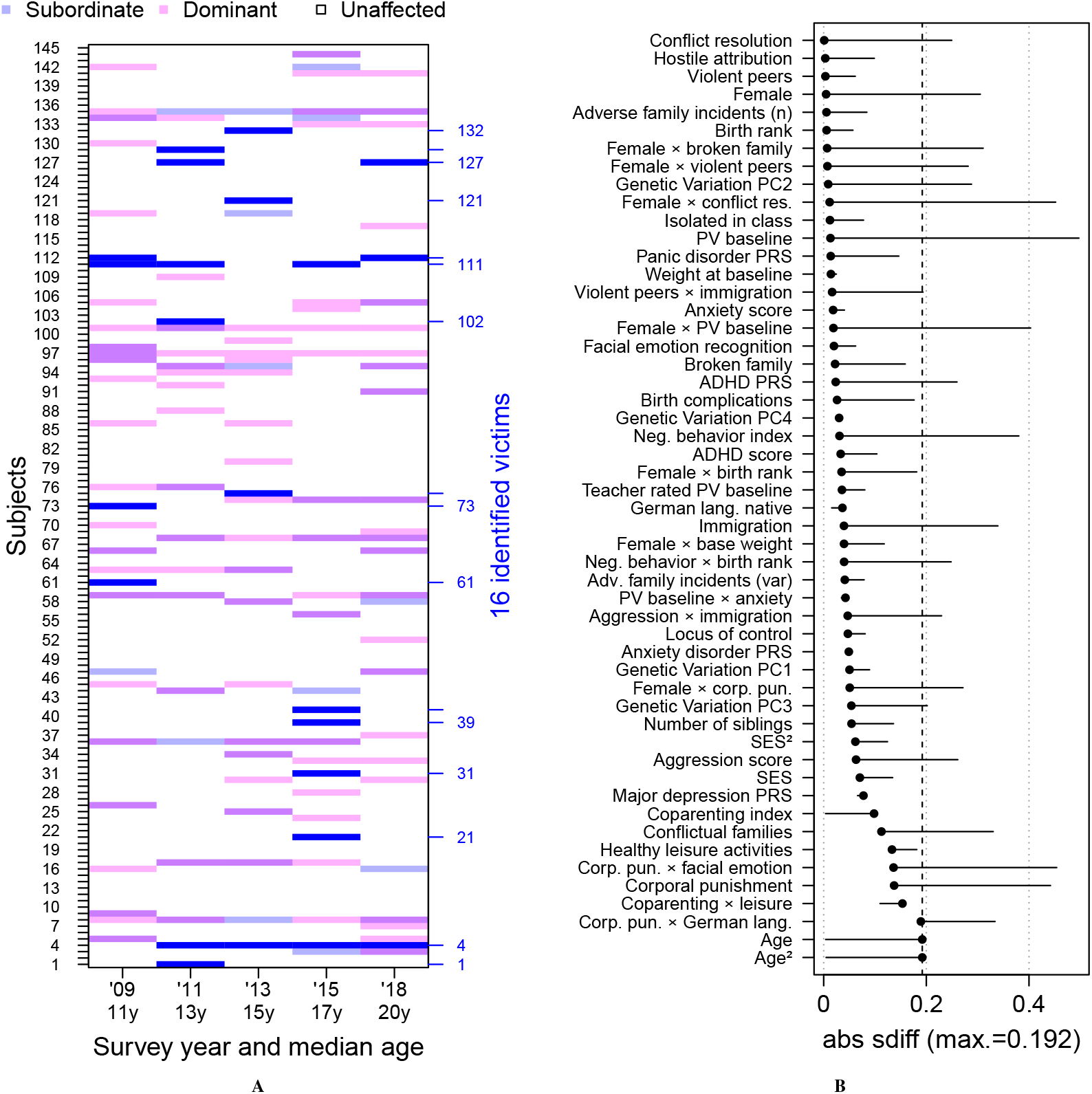
Peer victimization. (**A**) Prevalence of social peer adversity. The figure shows subordinate experiences and dominant behavior above 90% quantiles for each survey year. As the highest deciles, subjects above these thresholds are considered to be severely affected from the respective phenomenon. 16 individuals (7 male and 9 female) subjected to only subordinate experiences are identified as peer-victimized at the time of our study at around age 22. (**B**) Balancing after IP-weighting. The lollipop-plot shows absolute standardized differences between treated (PV) and control after IPW.

**Figure S2:**
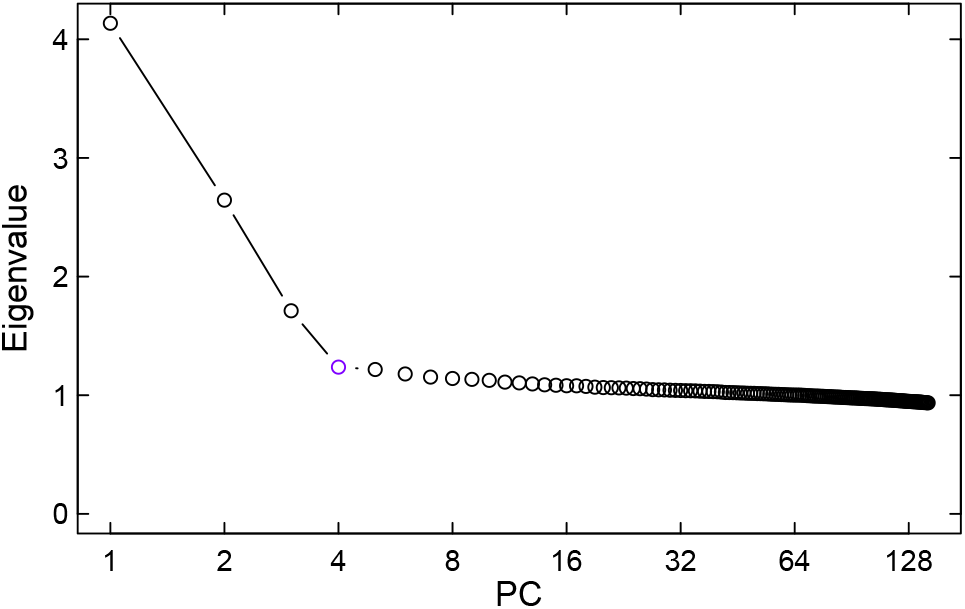
Scree plot of eigenvalues from genotyping, showing a clear inflection point at the fourth principal component.

**Figure S3:**
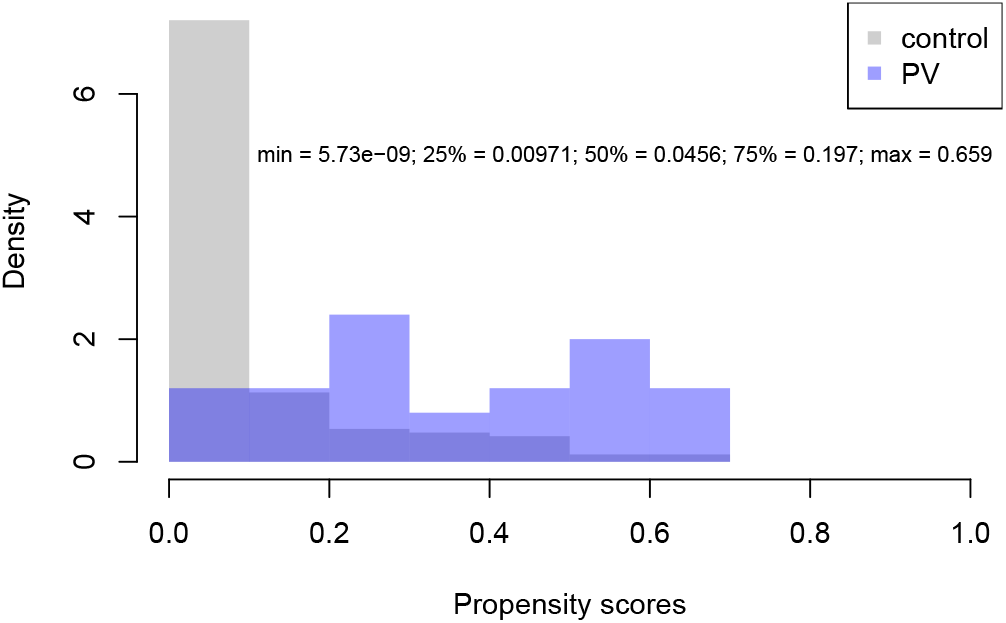
Propensity score distributions. Histogram of propensity scores for treated and control groups in the final analytic sample, indicating sufficient overlap for group comparability.

**Figure S4:**
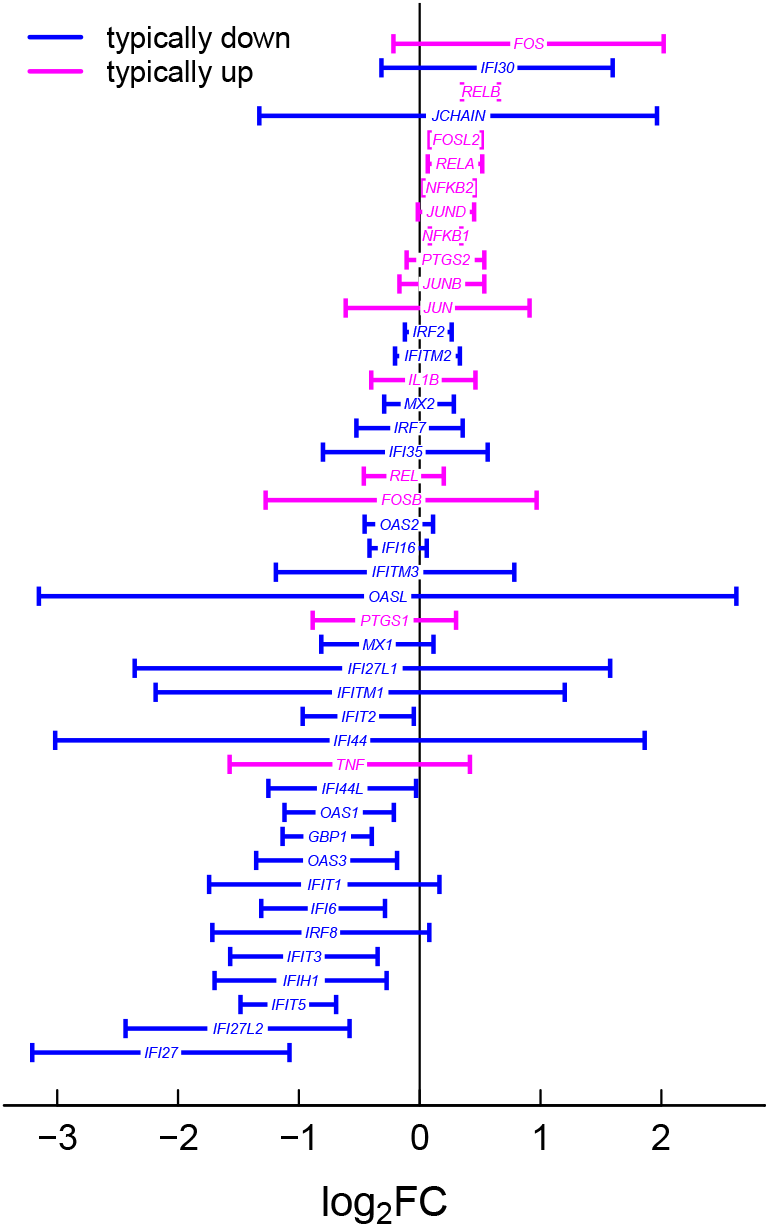
Up- and downregulation of CTRA genes in victims. Expression of 61 genes defining the Conserved Transcriptional Response to Adversity (CTRA). Genes typically upregulated in response to stress are shown in red, and those typically downregulated are shown in blue. Error bars represent 95% confidence intervals.

**Table S5:** Cytokines A—concentration profiles base model.

| | $\beta$ | SE | $t$ | $p$ | $p_{adj.}$ | 2.5% | 97.5% |
| --- | --- | --- | --- | --- | --- | --- | --- |
| *Pro-inflammatory:* |  |  |  |  |  |  |  |
| CCL11 | −0.034 | 0.174 | −0.196 | 0.8449 | 0.8811 | −0.378 | 0.310 |
| CCL4 | 0.122 | 0.050 | 2.448 | 0.0160 | 0.0511 | 0.023 | 0.221 |
| CCL5 | −0.173 | 0.185 | −0.940 | 0.3492 | 0.5609 | −0.540 | 0.193 |
| CXCL10 | −1.137 | 0.189 | −6.007 | <0.0001 | <0.0001 | −1.512 | −0.762 |
| CXCL9 | −1.123 | 0.215 | −5.229 | <0.0001 | <0.0001 | −1.549 | −0.697 |
| IL9 | −0.008 | 0.056 | −0.150 | 0.8811 | 0.8811 | −0.120 | 0.103 |
| LTA | 0.161 | 0.187 | 0.862 | 0.3908 | 0.5685 | −0.209 | 0.531 |
| TNF | 0.559 | 0.215 | 2.604 | 0.0105 | 0.0420 | 0.134 | 0.985 |
| *Anti-inflammatory:* |  |  |  |  |  |  |  |
| HGF | 0.058 | 0.186 | 0.312 | 0.7556 | 0.8811 | −0.311 | 0.427 |
| *Pleiotropic:* |  |  |  |  |  |  |  |
| CSF1 | −0.308 | 0.193 | −1.594 | 0.1138 | 0.2601 | −0.690 | 0.075 |
| CXCL12 | −0.150 | 0.106 | −1.704 | 0.0913 | 0.2435 | −0.391 | 0.030 |
| IL16 | −0.081 | 0.185 | −0.440 | 0.6605 | 0.8807 | −0.448 | 0.285 |
| *Other:* |  |  |  |  |  |  |  |
| CLEC11A | −3.270 | 0.818 | −4.000 | <0.0001 | 0.0006 | −4.891 | −1.649 |
| CCL27 | −0.043 | 0.228 | −0.188 | 0.8514 | 0.8811 | −0.496 | 0.410 |
| KITLG | −0.002 | 0.111 | −0.014 | 0.9885 | 0.9885 | −0.222 | 0.219 |
| PDGFB | 0.409 | 0.281 | 1.458 | 0.1477 | 0.2954 | −0.147 | 0.965 |

Comparative analysis of cytokine levels to characterize immune patterns in the investigated biological samples. Without leukocyte coding genes in the model.

**Table S6:** Cytokines B—concentration profiles LKC controlled model.

| | $\beta$ | SE | $t$ | $p$ | $p_{adj.}$ | 2.5% | 97.5% |
| --- | --- | --- | --- | --- | --- | --- | --- |
| *Pro-inflammatory:* |  |  |  |  |  |  |  |
| CCL11 | −0.024 | 0.142 | −0.171 | 0.8643 | 0.9220 | −0.306 | 0.257 |
| CCL4 | 0.131 | 0.042 | 3.143 | 0.0022 | 0.0087 | 0.049 | 0.215 |
| CCL5 | −0.350 | 0.172 | −2.034 | 0.0445 | 0.1185 | −0.692 | −0.009 |
| CXCL10 | −1.108 | 0.160 | −6.942 | <0.0001 | <0.0001 | −1.424 | −0.791 |
| CXCL9 | −1.096 | 0.220 | −4.982 | <0.0001 | <0.0001 | −1.532 | −0.660 |
| IL9 | 0.032 | 0.058 | 0.556 | 0.5791 | 0.8424 | −0.082 | 0.146 |
| LTA | 0.106 | 0.143 | 0.739 | 0.4616 | 0.7385 | −0.178 | 0.389 |
| TNF | 0.571 | 0.194 | 2.938 | 0.0041 | 0.0130 | 0.186 | 0.957 |
| *Anti-inflammatory:* |  |  |  |  |  |  |  |
| HGF | 0.034 | 0.187 | 0.180 | 0.8576 | 0.9220 | −0.338 | 0.405 |
| *Pleiotropic:* |  |  |  |  |  |  |  |
| CSF1 | −0.240 | 0.172 | −1.397 | 0.1654 | 0.2941 | −0.581 | 0.101 |
| CXCL12 | −0.150 | 0.090 | −1.675 | 0.0970 | 0.2218 | −0.327 | 0.028 |
| IL16 | 0.051 | 0.167 | 0.303 | 0.7622 | 0.9220 | −0.280 | 0.381 |
| *Other:* |  |  |  |  |  |  |  |
| CLEC11A | −3.116 | 0.656 | −4.753 | <0.0001 | <0.0001 | −4.416 | −1.816 |
| CCL27 | 0.061 | 0.191 | 0.321 | 0.7492 | 0.9220 | −0.317 | 0.440 |
| KITLG | −0.002 | 0.111 | −0.014 | 0.9885 | 0.9885 | −0.222 | 0.219 |
| PDGFB | 0.286 | 0.190 | 1.509 | 0.1344 | 0.2687 | −0.090 | 0.663 |

Comparative analysis of cytokine levels to characterize immune patterns in the investigated biological samples. With included leukocyte coding genes in the model.

